# Functional analysis of small-molecule and protein binding domains using monomeric streptavidin-based two-component biosensors

**DOI:** 10.64898/2026.11.03.763809

**Authors:** Roxane Mutschler, Zhong Guo, Maria Micaela Fiorito, Kirill Alexandrov, Zhenling Cui

## Abstract

Protein biosensors are important analytical and diagnostic tools that convert molecular recognition into measurable activity. Most common two-component biosensors require two non-competing recognition domains, yet binding affinity and specificity alone do not reliably predict their compatibility with the rest of the biosensor. We report a modular platform that exploits monomeric streptavidin (mSA2) as a surrogate recognition domain, enabling rapid functional evaluation of candidate binders in the context of biosensor scaffold without requiring a second ligand-specific binding protein. We established a chromatography-free protocol for recovering functional mSA2-based biosensor components from *Escherichia coli*. Using biotinylated small molecules methotrexate and vitamin D^3^, as well as heart failure protein biomarkers galectin-3 and S100A7 we demonstrate ligand-dependent reporter biosensor activation. The developed platform provides a scalable approach for rapid binder evaluation and biosensor prototyping, streamlining biosensor development using binding domains generated by directed evolution or computational protein design.

## Introduction

Protein-based biosensors are powerful tools for the detection and quantification of molecular targets in medical and veterinary diagnostics, food processing, environmental monitoring, and biomanufacturing. They function by converting binding of a ligand to a protein receptor into measurable activity. Two-component proximity-based biosensors have emerged as versatile and generalizable platforms for translating molecular recognition into measurable signals ^1–3^. These systems employ two ligand recognition domains that are coupled to sequences that form an active reporter when their local concentration rises due to the lig-and-mediated scaffolding (Figure 1A). For large targets, such as proteins, this can be achieved using two independently selected binders recognising distinct, non-over-lapping epitopes. In contrast, small molecules generally lack sufficient accessible surface area to accommodate two independent binders, requiring alternative recognition strategies such as chemically induced dimerisation (CID) ^4^. In these systems, a primary ligand binding domain, commonly termed the anchor, recruits a secondary recognition domain upon ligand binding, often through formation of a lig-and-dependent composite interface ^2, 4^.

**Figure 1:**
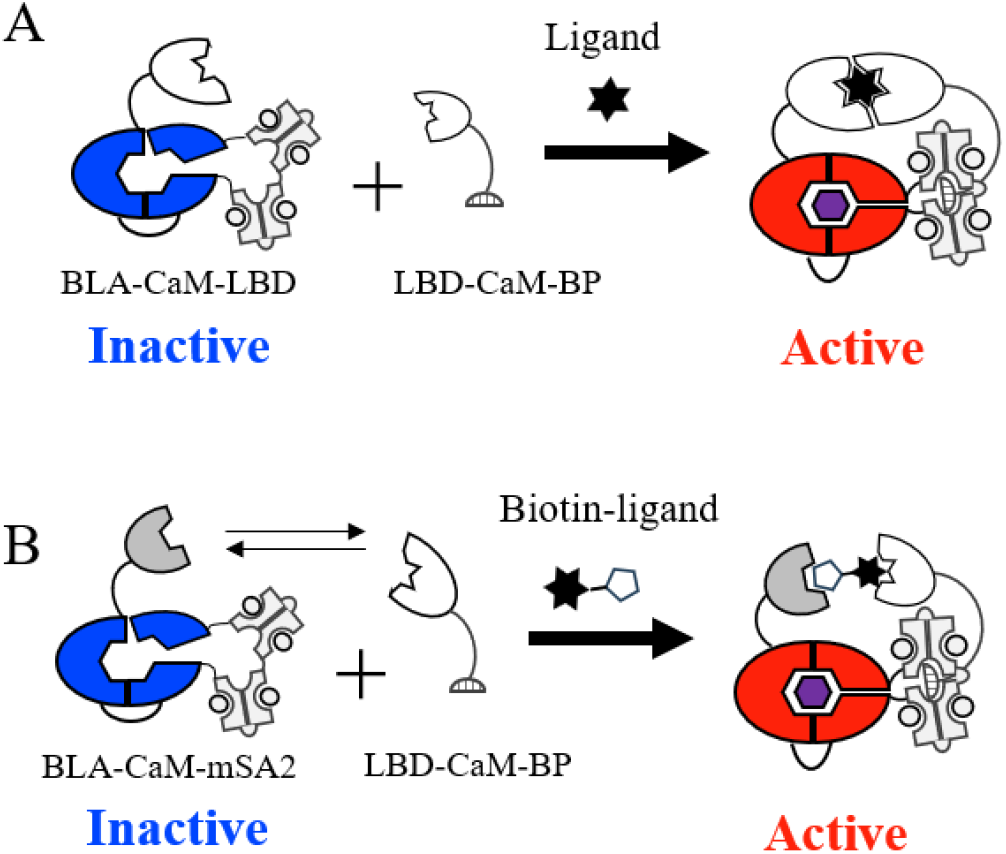
Monomeric streptavidin as surrogate receptor in two-component protein biosensors. **(A)** A schematic representation of two-component biosensor composed of a β-lactamase-calmodulin (BLA-CaM) chimera and a low affinity calmodulin-binding peptide (CaM-BP) fused to ligand-binding domain (LBD). Ligand recognition brings the two components into proximity, promoting CaM/CaM-BP association and BLA activation. **(B)** Adapted two-component biosensor using monomeric streptavidin (mSA2) as a universal biotin-binding anchor fused to either BLA-CaM or CaM-BP, with the candidate ligand-binding domain fused to the complementary component. Biotinylated ligand binding promotes ternary complex formation and BLA activation. Double arrows indicate interchangeable ligand-binding domains, enabling modular functional evaluation.

Advances in the AI-driven protein engineering and molecular display methods including phage and mRNA display, have enabled the generation of vast libraries of artificial protein receptors with tailored structures and ligand specificity ^4–7^. This creates a bottleneck in our ability to assess such receptors for affinity, selectivity and suitability for biosensor construction. The former properties are typically assessed using binding assays such as enzyme-linked im-munosorbent assays (ELISA), surface plasmon resonance (SPR) or biolayer interferometry (BLI). However, conventional binder selection and characterisation methods primarily evaluate binding affinity and specificity without assessing their compatibility with the reporter architecture. Consequently, even high-affinity binders may fail to generate robust biosensor responses, necessitating extensive downstream optimization ^3, 6^. The situation is particularly complicated with small molecules where development of CID further delays analysis of component’s compatibility.

Biotin-streptavidin interaction is widely used in the selection and characterization of artificial protein receptors ^8^. Biotinylated targets are commonly immobilized on streptavidin functionalised matrices for screening molecular display libraries and for molecular interaction characterization by methods such as ELISA, BLI and SPR ^4, 6^. The exceptionally strong interaction between biotin and streptavidin (K_D_ ~10^−14^ M) ^9^ provide a robust platform for target capture. However, the tetrameric structure of streptavidin and its propensity to misfold when expressed in prokaryotic hosts, complicates its use in modular biosensor systems ^10^. Protein sequence and evolutionary analysis suggested that tetramerisation is not strictly required for high-affinity biotin binding (Supplementary Figure 1), prompting the development of monovalent tetramers and monomeric streptavidin ^11^. Early monomeric variants suffered from poor biophysical stability and reduced affinity. The introduction of disulfide bonds and stabilising mutations addressed some of these stability issues, although its affinity for biotin (K_D_ ≈ 38 nM) remained substantially lower than that of wild type tetrameric streptavidin ^12^. Subsequent engineering incorporated structural elements from rhizavidin, a high-affinity biotin-binding protein that forms a dimer rather than the tetramers characteristic of avidin and streptavidin, yielding the mSA variant (PDB: 4JNJ) ^13^. To further reduce biotin dissociation, a histidine substitution at position 25 was introduced to create mSA2, providing additional van der Waals shielding of the biotin-binding pocket (K_D_ = 0.7 nM) ^14, 15^.

Here, we present a strategy for assessment of artificial lig- and binding domains for their compatibility with a two-component biosensor architecture ^6, 16^. We utilize the monomeric streptavidin (mSA2) as a surrogate second binder that recognises biotinylated ligands. Unlike conventional binding assays, this approach provides a functional readout that reflects both target recognition and the ability of receptor-reporter fusions to support signal generation. By linking receptor discovery to functional validation, this approach streamlines development of proximity-based protein biosensors.

## Results

### Development and validation of a modular mSA2-assisted proximity biosensor platform

To establish a platform for functional evaluation of ligand-binding domains, we adopted a previously developed two-component β-lactamase-calmodulin (BLA-CaM) chimeric biosensor (Figure 1A) ^6^. In this system, target-mediated interaction between BLA-CaM chimera and a calmodulin-binding peptide (CaM-BP) stimulates β-lactamase that can be quantified by an array of methods ^17^. To eliminate the requirement for a second target-specific recognition domain during initial functional screening, we employed monomeric streptavidin (mSA2) as a surrogate binding partner. mSA2 was fused to either BLA-CaM or CaM-BP, generating two complementary reporter configurations capable of recognising biotinylated targets (Figure 1B).

Consistent with the expression behavior of mSA2, both BLA-CaM-mSA2 and mSA2-CaM-BP were efficiently expressed in *Escherichia coli* accumulating predominantly in insoluble fractions (Supplementary Figure 2). We sought to establish a protein recovery strategy based on inclusion body solubilization and refolding ^12, 14, 15, 18^. Five combinations of solu-bilisation and refolding buffers were evaluated for each construct (Figure 2; Supplementary Table 1). Solubilisation buffers S2 and S3, combined with refolding buffer R2, yielded the highest recovery of soluble protein. This may be related to high arginine content of R2 buffer that has been previously identified as effective refolding additive. SDS-PAGE analysis revealed that both fusion proteins were isolated in high purity (~90%) without the need for affinity chromatography, providing an efficient route for producing the mSA2-based biosensor components.

**Figure 2:**
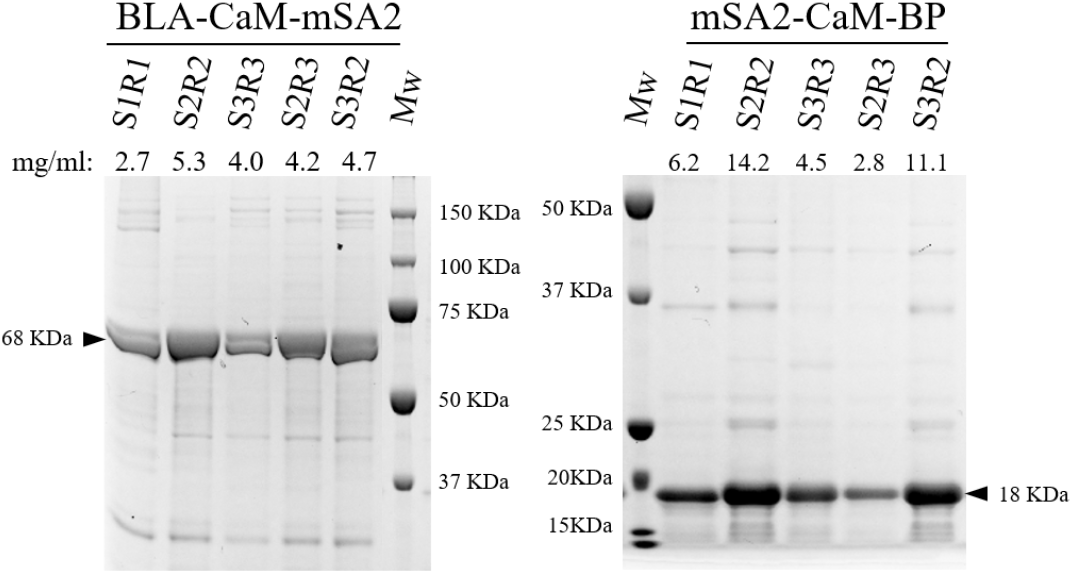
Purification and refolding of monomeric streptavidin-biosensor component fusion proteins. SDS-PAGE analysis of BLA-CaM-mSA2 (left) and mSA2-CaM-BP (right) recovered from inclusion bodies following denaturation and refolding under different combinations of solubilisation (S) and refolding (R) conditions: S1R1, S2R2, S3R3, S2R3 and S3R2. Protein concentrations (mg/ml), measured after refolding, are indicated above each lane.

### Functional characterization of mSA2-based biosensor components

To assess whether the refolded BLA-CaM-mSA2 fusion proteins were functional we tested their enzymatic activity in the presence of M13 calmodulin-binding peptide. Addition of M13 peptide to BLA-CaM-mSA2 increased enzymatic activity across the tested refolding conditions, with conditions S1R1, S2R2, and S3R3 displaying highest reaction rates (Figure 3A). These results demonstrate that the refolded BLA-CaM-mSA2 retained ligand dependent activation.

**Figure 3:**
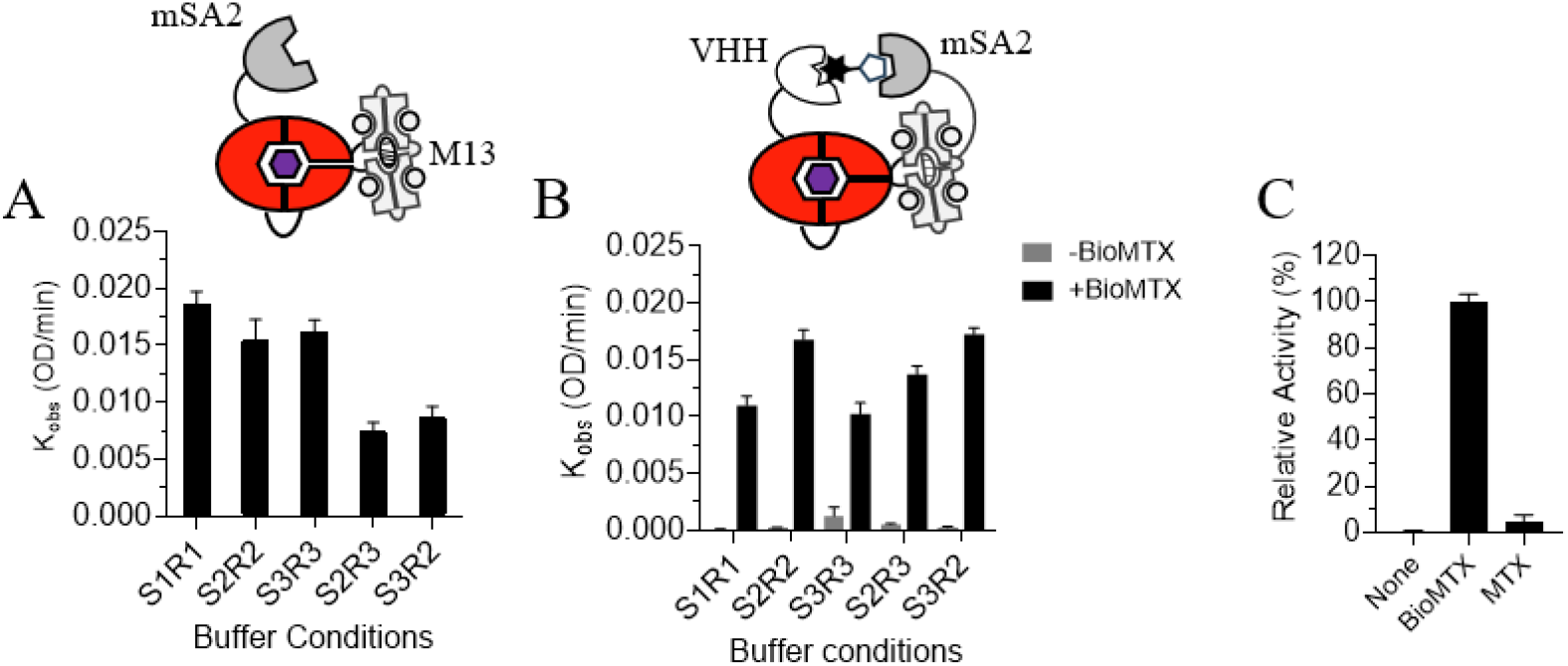
Functional evaluation of mSA2-based biosensor components prepared under different solubilisation and refolding conditions. **(A)** A plot of enzymatic activity of 25nM solution of BLA-CaM-mSA2 recovered under five different solubilisation (S) and refolding (R) conditions, assessed using a M13 peptide-mediated β-lactamase activity in the presence or absence of 200 nM M13 peptide. The reaction was incubated for 15 min in buffer containing 20 mM Tris-HCl (pH 7.2), 100 mM NaCl and 1 mM CaCl□β-lactamase activity was quantified by adding 50 µM UW154 substrate, and recording absorbance at 520 nm for 30 min. The initial reaction rates were calculated from the linear slopes of the absorbance traces. **(B)** Plot of initial rates of 150nM solution of mSA2-CaM-BP recovered under the same five solubilisation and refolding conditions as in (A) and mixed with 25 nM BLA-CaM-VHH in the presence or absence of 500 nM biotinylated methotrexate (BioMTX). Measured as in (A). **(C)** A plot of normalized activity of a two-component biosensor comprising BLA-CaM-VHH and mSA2-CaM-BP (purified at preparative scale) in the presence or absence of 500 nM biotinylated methotrexate (BioMTX) or unmodified methotrexate (MTX). Enzymatic activity of all reactions was normalised to the BioMTX-induced response. β-Lactamase activity was measured as described in A. Data are presented as mean ± s.d. of three independent measurements.

To assess the functionality of refolded mSA2-CaM-BP, we adopted a previously developed two-component metho-trexate (MTX) biosensor ^19^. It comprises an MTX-binding VHH domain fused to BLA-CaM (BLA-CaM-VHH), and a nanoCLAMP domain fused to CaM-BP (nanoCLAMP-CaM-BP). The nanoCLAMP specifically recognized the MTX-bound VHH complex, bringing the two components into proximity and activating β-lactamase. To evaluate mSA2 as a universal anchor domain, we replaced nanoCLAMP-CaM-BP with mSA2-CaM-BP while retaining BLA-CaM-VHH as the ligand-recognition component. The resulting BLA-CaM-VHH/mSA2-CaM-BP system exhibited ligand-dependent enzymatic activation in the presence of biotinylated MTX (BioMTX) (Figure 3B). Protein refolded under S2R2 and S3R2 conditions displayed highest reaction rates. These findings demonstrate that the refolded mSA2-CaM-BP retained the functionality required for its biosensor activation.

Based on soluble mSA2 fusion proteins and their activity analysis, the S2R2 condition was selected for subsequent large-scale biosensor component production (Figure 2 and 3A, B). The functionality of the proteins purified on prepar-ative scale was further evaluated. We confirmed that the mSA2-CaM-BP-based biosensor exhibited robust activation in the presence of BioMTX, whereas unmodified MTX induced minimal activity (Figure 3C). These results demon-strate that mSA2 enables proximity-dependent reporter activation without second ligand-specific binder.

### Analysis of small molecule and protein binding domains using mSA2-based biosensor

Next, we investigated the utility of the developed platform in analysis of ligand binding domains. As a first example we chose artificial Vitamin-D_3_ binding protein CDL2.3a ^20^. The protein was fused to BLA-CaM and CaM-BP and paired with the complementary mSA2 fusion proteins in a reaction mixture. Their activity was evaluated in the presence or absence of a biotinylated Vitamin D_3_ conjugate. Both systems displayed ligand dependent β-lactamase activation (Figure 4A and 4B), demonstrating that mSA2-assited proximity sensing can be extended to different small-molecule recognition domains.

**Figure 4.**
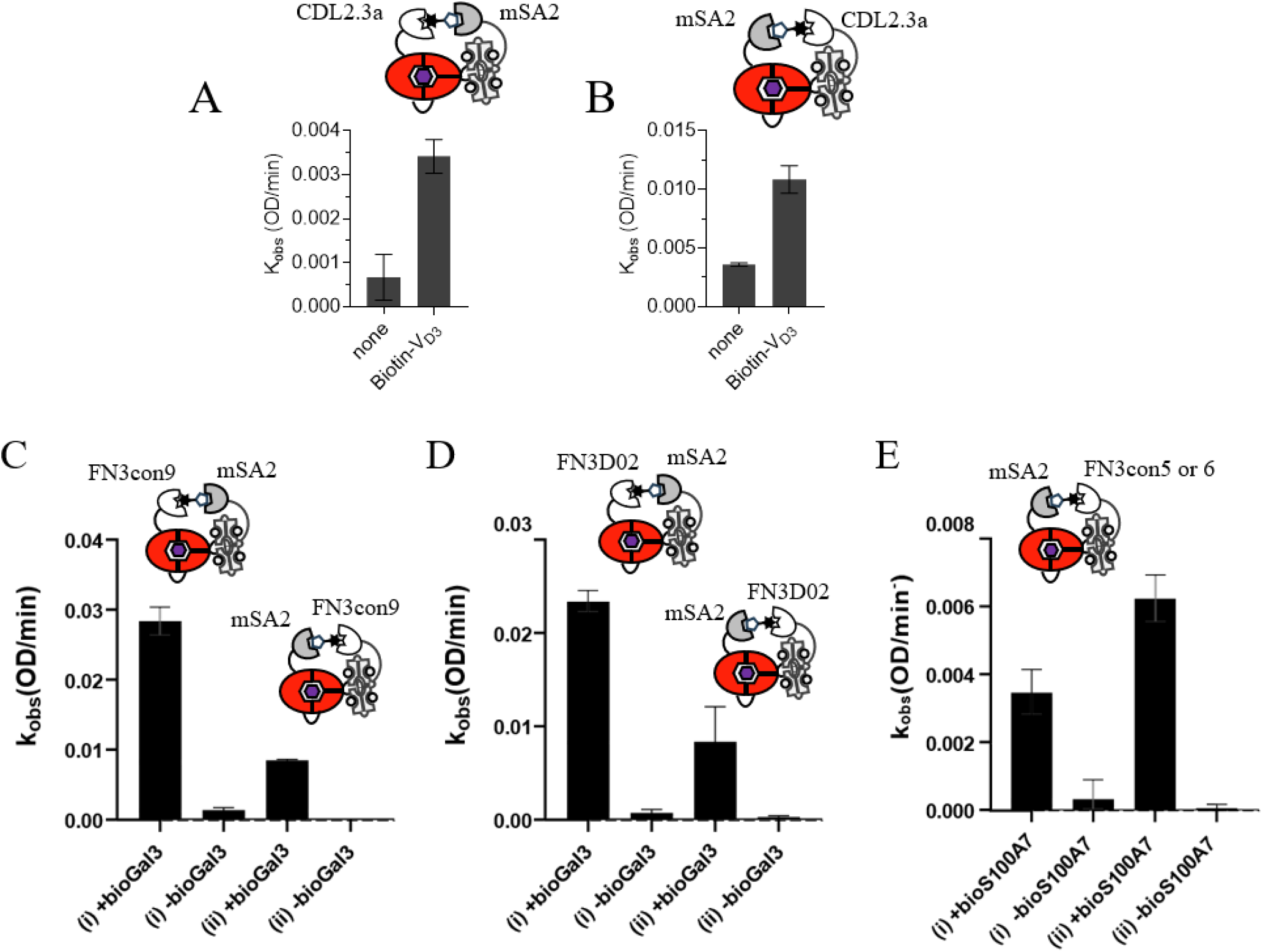
Functional evaluation of small-molecule and protein binding domains using mSA2-based two-component biosensors. **(A, B)** Enzymatic activity of biosensors comprising the vitamin D_3_-binding domain CDL2.3a and mSA2 in two alternative configurations: **(A)** BLA-CaM-CDL2.3a (25 nM) paired with mSA2-CaM-BP (150 nM) and **(B)** BLA-CaM-mSA2 (25 nM) paired with CDL2.3a-CaM-BP (100 nM), in the presence or absence of biotinylated vitamin D_3_ (Biotin-V_D3_). **(C, D)** Enzymatic activity of biosensors comprising the galectin-3-binding domains FN3con9 **(C)** or FN3D02 **(D)**, paired with mSA2 in two alternative configurations: (i) BLA-CaM-LBD (25 nM) paired with mSA2-CaM-BP (150 nM) and (ii) BLA-CaM-mSA2 (50 nM) paired with LBD-CaM-BP (100 nM), in the presence or absence of 500 nM biotinylated galectin-3 (bioGal3). **(E)** Enzymatic activity of biosensors comprising BLA-CaM-mSA2 (50 nM) paired with the S100A7-binding domains FN3con5 or FN3con6 fused to CaM-BP (150 nM), in the presence or absence of biotinylated S100A7 (bioS100A7). β-Lactamase activity was measured as described in Figure 3A. Data are presented as mean ± s.d. of three independent measurements.

Next, we wanted to test whether the developed biosensor system could be used to analyse binding domains selected to protein targets. To this end, we utilized FN3con domains that were previously selected using mRNA display against the heart failure-associated protein biomarkers Galectin-3 and S100A7 ^1, 3^. Open reading frames of the Galectin-3 binding domains, FN3D02 and FN3con9, were cloned into both BLA-CaM and CaM-BP constructs. The resulting fusion proteins were expressed in *Escherichia coli* and purified to homogeneity. We then tested activity of these binder fusion proteins paired with the complementary mSA2 fusion, mSA2-BLA-CaM or mSA2-CaM-BP, in the presence or absence of biotinylated Galectin-3 or S100A7. In both cases we recorded target-dependent activation of biosensor’s activity (Figure 4C, D and E).

## Discussion

The functional characterisation of ligand-binding proteins remains a technical bottleneck of protein engineering, particularly when evaluating large numbers of candidates generated through directed evolution or computational design. Although binding assays can establish target recognition and affinity, translating these interactions into functional biosensors often requires additional engineering, including the identification of a second non-competing binding domain. Here, we developed a modular two-component biosensor platform that uses monomeric streptavidin (mSA2) as a universal biotin-binding domain, enabling lig-and-dependent enzymatic activation through the interaction of a candidate binding protein with its biotinylated target. By replacing one target-specific recognition domain with mSA2, this approach simplifies the assembly of functional biosensors and provides a versatile strategy for evaluating ligand-binding domains.

A key feature of this platform is its compatibility with two alternative fusion configurations, in which mSA2 can be coupled to either the allosteric β-lactamase reporter or to the calmodulin-binding peptide (CaM-BP). We established a chromatography-free purification procedures for recovering both fusion proteins from inclusion bodies and systematically compared solubilisation and refolding conditions based on soluble protein recovery and functional activity. The S2R2 condition provided a suitable balance between protein recovery and activity for both fusion proteins and was selected for subsequent large-scale production.

We demonstrated the applicability of the platform using a vitamin D_3_-binding protein, CDL2.3a, and binders targeting the protein biomarkers galectin-3 and S100A7. Ligand-dependent activation was observed across these distinct targets, supporting the adaptability of the mSA2-assisted architecture to both small molecules and proteins. The use of different binding scaffolds further suggests that the platform is not restricted to a particular receptor architecture, provided that the target can be biotinylated without disrupting the relevant binding interaction. These findings establish the potential of mSA2-assisted biosensors as a generalisable approach for functional assessment of ligand-binding proteins.

Future work will focus on extending the platform to high-throughput screening of candidate binders. By integrating the mSA2-assisted biosensor assay with *in vitro* protein production, large number of candidate binders could be evaluated for their ability to function in the context of a protein biosensor. This approach facilitates identification of functionally compatible binders at an early stage of biosensor development, reducing attrition at the later stages of biosensor design.

## Acknowledgements

This work was partially funded by the ARC Centre of Excellence in Synthetic Biology CE200100029, ARC Linkage Project LP200200916, and NHMRC Investigator grant APP2033951 to KA. This work was also supported by a QUT-CSIRO Fellowship awarded to ZC (QUT PURE Research ID: 178477198). KA and ZC gratefully acknowledge the financial support of CSIRO-QUT Synthetic Biology Alliance as well as the support of CSIRO Synthetic Biology and Advanced Engineering Biology Future Science Platforms.

## Author contributions

R.M. contributed to validation, formal analysis, investigation, data curation, visualization, and writing the original draft. Z.G. contributed to conceptualization and methodology. M.F. contributed to resources and manuscript review. K.A. contributed to supervision, funding acquisition, and manuscript review and editing. Z.C. contributed to conceptualization, methodology, investigation, data curation, supervision, and manuscript review and editing.

## Competing interest statement

No conflict of interest.

## Notes

### Competing Interest Statement

The authors have declared no competing interest.

