## Supplemental Files for "Functional analysis of small-molecule and protein binding domains using monomeric streptavidin-based two-component biosensors"

Supplementary Materials for

**A versatile monomeric streptavidin-based platform for functional screening of small molecule and protein binders**

### The file includes:

Materials and Methods

Supplementary Figures 1-3

Supplementary Tables 1-2

References

### Materials and Methods

**Design and Construction of Recombinant Expression Plasmids**

The DNA and amino acid sequences of the recombinant proteins, including the β-lactamase–calmodulin (BLA-CaM) switch and the low-affinity calmodulin-binding peptide (CaM-BP), are provided in Supplementary Tables 2 and 3. The corresponding coding sequences were synthesised by Gene Universal and cloned into the pET28a expression vector using NcoI and HindIII restriction sites, as previously described ^1^. To generate the monomeric streptavidin (mSA2)-based biosensor constructs, the mSA2 coding sequence was inserted into the corresponding BLA-CaM and CaM-BP expression vectors using BamHI and SalI restriction sites, yielding BLA-CaM-mSA2 and mSA2-CaM-BP, respectively. Similarly, receptor-fused biosensor constructs (BLA-CaM-receptor and receptor-CaM-BP) were generated by inserting the respective ligand-binding domain sequences using the same restriction sites. Specifically, DNA fragments encoding the methotrexate binding VHH domain and Vitamin D_3_ binding domain (CDL2.3a) were cloned to generate BLA-CaM-VHH, VHH-CaM-BP, BLA-CaM-CDL2.3a and CDL2.3a-CaM-BP. All constructs were generated by Gene Universal and verified by DNA sequencing. The galectin-3 binding domains, FN3con9 and FN3D02, and the S100A7 binding domains FN3con5 and FN3con6 were incorporated into BLA-CaM and/or CaM-BP based biosensor constructs as described in our previous studies ^2, 3^. The corresponding recombinant expression plasmids were used in this study.

**Expression of monomeric streptavidin constructs**

Sequence-verified plasmids encoding BLA-CaM-mSA2 and mSA2-CaM-BP were transformed into *Escherichia coli* BL21(DE3) cells and plated on LB agar supplemented with kanamycin, followed by overnight incubation at 37 °C. A single colony of each construct was inoculated into 1.6 mL LB broth containing kanamycin and cultured overnight at 37 °C. The overnight cultures were subsequently used to inoculate three and two 1L LB cultures for BLA-CaM-mSA2 and mSA2-CaM-BP, respectively. Cells were grown at 37 °C until the optical density at 600 nm reached approximately 0.8, at which point protein expression was induced with 0.3 mM isopropyl β-D-1-thiogalactopyranoside (IPTG). Cultures were incubated overnight at 18 °C, and cells were harvested by centrifugation at 4,000 rpm for 12 min, yielding approximately 22 g and 7 g of wet cell pellets for BLA-CaM-mSA2 and mSA2-CaM-BP, respectively.

**Inclusion body isolation, solubilisation and refolding optimisation**

Three litres of BLA-CaM-mSA2 cell culture yielded 22 g of cell pellet, from which 0.65 g was allocated for small-scale optimisation, while the remainder was stored at −80 °C. Similarly, two litres of mSA2-CaM-BP culture produced 7 g of cell pellet, of which 0.45 g was used for small-scale optimization and the remainder stored at -80 °C.

Frozen cell pellets were thawed at room temperature (22 °C), centrifuged at 23,000 g at 4 °C for 15 minutes and resuspended in 2 ml of disruption buffer containing 1X BugBuster protein extraction reagent (Merck, #70921), 0.1 mg/ml DNAaseI and 20 µg/ml lysozyme. Following incubation at 22 °C for 20 minutes, samples were centrifuged at 5,000 g for 15 minutes at 4 °C. The resulting pellets were washed three times with 1.5 ml of washing buffer (30 mM Tris-HCl, pH 8.0, 2 mM EDTA and 0.1% Triton X-100) and centrifugation at 15,000 g for 5 minutes at 4 °C between washes.

The washed inclusion body pellets were resuspended in 1 ml of solubilisation buffer and divided into three aliquots for evaluation under different solubilization conditions (S1-S3). Following solubilization, samples were centrifuged at 5,000 g for 5 minutes at 4 °C and then dialysed overnight at 4 °C against 2ml of refolding buffer corresponding to the solubilisation buffer used (e.g., S1 with R1) using a 96 well micro-dialysis plate (10KDa cutoff, #A50469, Thermofisher Scientific).

Following dialysis, the refolded protein solutions were collected into PCR tubes. Visible precipitation was observed in some samples, which were subsequently centrifuged at 23,000 g for 15 minutes at 4 °C to remove insoluble material. Protein concentrations in the clarified supernatants were measured by nanodrop using the respective refolding buffer as a blank. Meanwhile, 4 µl from each sample was reserved for SDS-PAGE analysis.

**Large scale purification of monomeric streptavidin constructs**

The remaining BLA-CaM-mSA2 (~21 g) and mSA2-BP (~6 g) cell pellets were resuspended in ∼66 ml and ∼35 ml of ice-cold resuspension buffer (50mM Tris-HCl pH8, 1mM EDTA, 0.1 mg/ml DNaseI and 0.1 mM AEBSF), respectively. Following incubation on ice for 20 minutes, cells were disrupted by two passes through high pressure cell disruptor (Constant Systems) at 26 kpsi. Cell lysates were centrifuged at 20,000 g for 30 minutes at 4 °C, and the resulting pellets were washed three times with 50 ml of ice-cold washing buffer (30 mM Tris-HCl pH 8, 2 mM EDTA, 0.1% Triton X-100). The first wash was followed by centrifugation at 20,000 g for 15 minutes at 4 °C whereas the second and third washes were centrifuged at 5,000 g for 15 minutes at 4 °C.

The inclusion bodies were then dissolved in 30 ml of solubilisation buffer S2, supplemented with 10 mM DTT. The solubilized proteins were subsequently dialysed three times against 200 ml of solubilisation buffer S2 for at least 4 hours each, followed by two rounds of dialysis against 250 ml of refolding buffer R2 for at least 4 hours each. Finally, the protein was dialysed overnight against 2 litres of storage buffer (20 mM Tris-HCl, pH 7.2, 100 mM NaCl). All dialysis steps were performed at 4 °C with continuous stirring at 200 rpm. Following dialysis, insoluble aggregates were removed by centrifugation at 5,000 g for 5 minutes at 4 °C. The clarified protein solutions were aliquoted and stored at -80 °C.

**Molecular cloning, expression and purification of biotinylated Galectin-3 and S100A7**

DNA fragments encoding Galectin-3 and S100A7, each incorporating an N-terminal His-tag and Avi-tag, were cloned into the pET28a plasmid using the NcoI and HindIII restriction enzymes by Gene Universal as described previously^2, 3^. The sequence verified plasmids were co-transformed with the BirA plasmid into BL21 (DE3) cells and plated onto LB agar plates containing kanamycin and chloramphenicol. Following overnight incubation at 37^o^C, colonies resuspended into 12 mL of LB broth. The resulting cell suspension was distributed into six 1L flasks containing LB broth supplemented with kanamycin and chloramphenicol and cultured at 37 ^o^C until the optical density at OD_600_ reached 0.8. Protein expression and *in vivo* biotinylation were induced by adding 0.3 mM IPTG and 25 mg/L of biotin. The cells were grown at 18 ^o^C overnight and then harvested by centrifugation at 4000 rpm for 12 minutes.

Cell pellets were resuspended in lysis buffer containing 300 mM sodium chloride, 50 mM sodium phosphate, 20 mM imidazole, 1 mM DNase1 and 1 mM AEBSF, pH 8. Cells were disrupted using a CFII high-pressure cell disrupter (Constant Systems, UK) at 27 kPsi, and lysates were clarified by centrifuged at 15000 rpm for 35 minutes. The resulting supernatant was collected and loaded onto a Ni-NTA His-Trap FF crude column using an ÄKTAexpress chromatography system. The column was equilibrated and washed with binding buffer (50 mM sodium phosphate, 300 mM NaCl, and 20 mM imidazole pH8.0) and bound proteins were eluted using elution buffer containing 50 mM sodium phosphate, 300 mM NaCl, and 500 mM imidazole pH 8.0.

Eluted proteins were concentrated using Amicon Ultra-15 Centrifugal Filter Unit (10 kDa MWCO) and further purified by size exclusion chromatography on a HiLoad 16/600 Superdex 75 pg column (Cytiva), pre-equilibrated with storage buffer (20 mM Tris-HCl, 100 mM NaCl, pH 7.2). Fractions corresponding to the expected protein elution peaks were collected, concentrated and subsequently aliquoted and stored at -80^o^C.

**Expression and purification of biosensor components**

Plasmids encoding the BLA-CaM and CaM-BP-based biosensor components were transformed into BL21 (DE3) competent cells (Thermo Fisher Scientific). Transformed cells were then cultured in LB broth supplemented with 50 µg/mL of Kanamycin and grown at 37 ^o^C until the optical density (OD_600_) reached 0.8, at which point 0.3 mM of IPTG was added for protein induction, followed by overnight incubation at 18 ^o^C. Recombinant proteins were purified using the nickel-affinity chromatography protocol described above for biotinylated galectin-3 and S100A7, omitting the size-exclusion chromatography step.

**Functional evaluation of binding domains using activity assays**

The functionality of refolded BLA-CaM-mSA2 proteins were initially assessed using a M13 calmodulin-binding peptide mediated β-lactamase activity assay. Reactions contained 25 nM BLA-CaM-mSA2 in assay buffer comprising 20 mM Tris-HCl (pH 7.2) and 100mM NaCl, 1mM CaCl_2_, in the presence or absence of 200 nM M13 peptide. β-lactamase activity was initiated by adding the chromogenic substrate UW154 to a final concentration of 50 μM, and absorbance was monitored at 520 nm using a BioTek Synergy Neo2 Reader.

Two component BLA-based biosensor assays were conducted under identical assay conditions, with BLA-CaM-mSA2 or BLA-CaM-receptor at 25 nM or 50 nM and mSA2-CaM-BP or receptor-CaM-BP at 100 nM or 150 nM in the presence or absence of 200 nM or 500 nM of biotinylated analyte. Biosensor kinetics were recorded after the addition of UW154, following the previously described protocol.

### Supplementary Figures

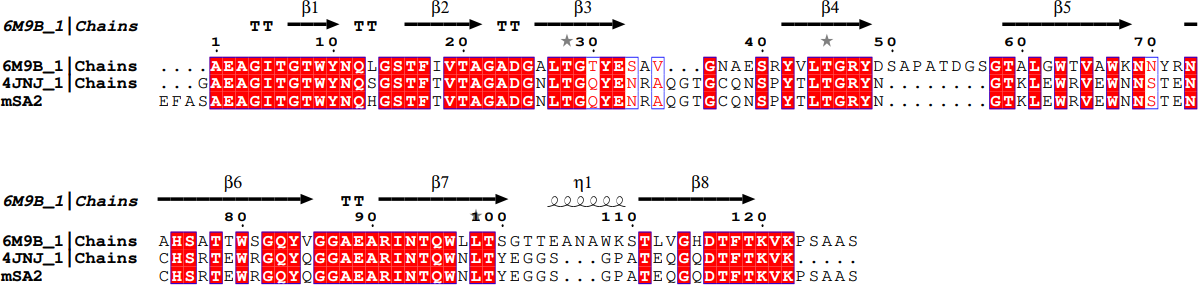

**Supplementary Figure 1. Sequence alignment of wild-type and engineered Streptavidin variants.** Multiple sequence alignment of wild-type tetrameric streptavidin (PDB: 6M9B), the first-generation monomeric streptavidin-rhizavidin hybrid (PDB: 4JNJ), and the second-generation engineered monomeric streptavidin (mSA2), generated using ESPript 3.0 ^4^. Identical residues across all three sequences are highlighted in red, whereas residues conserved between two sequences are indicated by red lettering with a blue outline. The alignment highlights conserved and variable regions among streptavidin variants, including residues potentially involved in structural stability and biotin recognition.

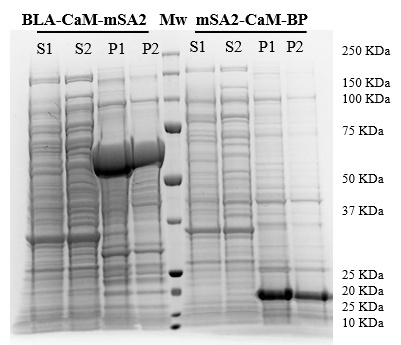

**Supplementary Figure 2. SDS-PAGE analysis of BLA-CaM-mSA2 and mSA2-CaM-BP expression and solubility**. Coomassie-stained SDS-PAGE gel showing the soluble and insoluble fractions of BLA-CaM-mSA2 and mSA2-CaM-BP expressed in *E. coli*. S1 and S2 represent the supernatant fractions following cell disruption, prepared without and with dithiothreitol (DTT), respectively. P1 represents the insoluble pellet fraction following cell disruption, whereas P2 represents the pellet after a single wash with phosphate-buffered saline (PBS). Equal volumes of supernatant and pellet fractions were loaded for comparison. Molecular weight markers (Mw) are indicated on the right.

**Supplementary Table 1.** Composition of buffers used for solubilisation and refolding of mSA2-fused biosensor proteins

| **Purification Buffers** |  |
| --- | --- |
| Disruption Buffer | 1X BugBuster protein extraction reagent (Merck, #70921, 1: 10 dilution), supplemented with 0.1 mg/ml DNAaseI, 20 µg/ml lysozyme |
| Washing Buffer | 30mM Tris-HCl (pH 8.0), 2 mM EDTA, 0.1% Triton X-100 |
| Solubilisation Buffer 1 (S1) | 6 M GdmCl (pH 1.5) |
| Solubilisation Buffer 2 (S2) | 6 M GdmCl, 50 mM Tris, 100 mM NaCl, 10 mM EDTA, 10 mM DTT, 5% glycerol |
| Solubilisation Buffer 3 (S3) | 6 M GdmCl, 50 mM Tris-HCl, 100mM NaCl, (pH 8.0) |
| Refolding Buffer 1 (R1) | 0.2 M NaHCO_3_ (pH 8.7) |
| Refolding Buffer 2 (R2) | 100 mM Tris, 400 mM Arg-HCl, 2 mM EDTA, 5 mM GSH, 0.5 mM GSSG, pH 8.0 |
| Refolding Buffer 3 (R3) | 50 mM Tris-HCl (pH 8.0), 150 mM NaCl, 0.2 mg/ml GSSG, 1 mg/ml GSH |
| Resuspension Buffer | 50mM Tris-HCl (pH 8.0), 1mM EDTA, 0.1 mg/ml DNaseI and 0.1 mM AEBSF |
| Storage Buffer | 20 mM Tris-HCl (pH 7.2), 100 mM NaCl |

**Supplementary Table 2.** Sequences of oligonucleotides used in this study.

| **mSA2** | GAGTTTGCGTCGGCGGAGGCGGGAATTACGGGCACTTGGTATAACCAA CATGGTTCGACCTTCACGGTTACCGCCGGGGCCGACGGGAATCTTACCG GGCAATACGAAAATCGTGCTCAAGGCACAGGGTGTCAAAACTCGCCATA CACGTTGACGGGGCGCTACAACGGGACAAAGCTGGAATGGCGTGTAGA ATGGAATAATTCGACCGAGAACTGCCACAGCCGTACTGAGTGGCGCGGA CAATATCAAGGGGGTGCGGAGGCTCGCATTAACACCCAATGGAACCTGA CCTATGAGGGAGGATCAGGTCCAGCCACCGAACAAGGGCAAGATACTTT  CACTAAAGTCAAACCCAGCGCTGCTAGT |
| --- | --- |
| **VHH_MTX_** | GGGTCTCAAGTGCAACTTGTCGAGAGCGGAGGCGGCTTAGTACAAGCCG GCGGGAGTTTGCGCTTTCTTGTGCTGCATCGCGCCGTTCAAGCCGCAGTT GGGCGATGGCATGGTTTCGCCAAGCGCCTGGGAAAGAACGTGAGTTCGT GGCAAAAATCTCTGGAGATGGCCGTTTAACGACTTATGGGGACTCAGTAA AGGGGCGTTTTACTATCTCACGCGACAATGCAGAGTATTTAGTATACTTGC AAATGGACAGTCTTAAGCCCGAAGACACAGCGGTCTACTATTGTGCCGCC GACGACAACTATGTTACTGCAAGCTGGCGTAGTGGCCCTGACTATTGGGG  GCAGGGGACGCAGGTCACAGTATCTTCT |
| **CDL2.3a (VD_3_**  **binding domain)** | TTGCCGACAGCCCATGAGGCCATCGAAGCCGCTCTTGCAGATTTCGTAAAG GTATACAACTCCAAAGACGCCGCTGGGGTCGCGTCTAAATACATGGACGAT GCTGTGATCTTTCCGCTGGATATGGCCCGCGTCGACGGTCGCCAAAACATT CAAAAGTTATGGCAGGGTTTAATGGATATGGGTGTGTCCGAACCAAAATTT ACTACACTGAATGTCCAAGAATCGGGCGACTTCGCGTTTGAATCAGGTTCG TTCTCCTTAAAGGGACCCGGCAAGGACTCCAAGTTGGTTGACATTGCTGGC ATCTACGTGGAGGTGTGGCGCAAGGGGCAAGACGGTGGGTGGAAGTTGT ACCGTACTATCGCAAATCTTGATCCAGCTCGTTTAGAACACCACCACCATCA  TCAC |
| **FN3con9 (Gal3 binding domain)** | CCATCGCCCCCAGGCAATTTGCGCGTTACAGATGTAACAAGCACTTCTGTCA CATTATCATGGGATGCGTACTCCGAATACTCACGCGGTTATCGTGTGGAGTAC CGTGAAGCCGGAGGGGAGTGGAAGGAAGTTACCGTTCATTTATTTGAGCA GTTCTACACAGTGACCGGCTTGAAACCGGGCACCGAATATGAGTTCCGCGT TCGTGCCGTGATTGGTTG GGATTACTGGATTTTCTCTTCGGTGTCTGTCACC ACC |
| **FN3D02**  **(Gal3 binding domain)** | ATGCGCGTTAGCGATGTTCGCGTGATTTGGAAGTCGTAGCTGCCACGCCAA CTTCCTTGCTTATTTCCTGGGATGCATCTAGTGTTTCAGTAGATTTTTACAATA TCACTTACGGTGAAACCGGGACGAACAGTCCGGTTCAAGCATTCAAGGTTC CAGGCTCGAAGAGCACAGCAACGATCAGTGGACTGAAGCCGGGTGTTGA  CTATACTATTACCGTCTACGGGTCCTACCCCTATCCTAAGCACATGACTACGG ATAATCCTATCTCCATCAATTATC GTACT |
| **FN3con5 (S100A7**  **binding domain)** | CCTAGTCCGCCGGGTAATTTGCGTGTAACCGATGTCACATCTACGAGTGTG ACCCTTTCATGGTGGCCAGAAATCGTCCCTTTGCACGGCTATCGTGTAGAG TACCGTGAGGCTGGGGGAGAATGGAAGGAGGTGACTGTGATCGACCGTG CTCCTGAATACACAGTCACCGGATTAAAGCCTGGCACGGAGTATGAATTCC GTGTACGCGCGGTGTCAGGCGCGTTATACTTCAACGCGTCATCAGTCTCTG  TTACTACT |
| **FN3con6 (S100A7**  **binding domain)** | CCAAGTCCTCCAGGAAATCTGCGCGTCACGGACGTAACTTCGACAAGTGTCA CACTGTCGTGGAATGACGAGTTCCGCGCGTGGCATGGGTACCGTGTTGAATA CCGCGAGGCTGGTGGTGAGTGGAAAGAGGTCACAGTAATTGAGTGGCCTAA GATTTACACAGTAACTGGGTTGAAGCCAGGTACGGAGTACGAGTTTCGTGTC CGTGCTGTCCAAGCTCGCTGGGAATGGACAACATCGAGCGTGTCGGTTACAA CG |
| **His-FLAG-**  **mSA2/ binding domain-**  **CaM-BP** | ATGGATAGCCATCATCATCACCATCACGGATCCGACTACAAAGATGACGATGA CAAAGGATCC—mSA2 / BINDING-DOMAIN—GTCGACGGCGGAAGTGGCTCC GGTGGTTCAGGTGGCGGTAGCGGTAGCTCCGGAGGAAGTGGCGGATCTGGG GGTGGAAAGCGTCGTTGGAAAAAGAACTTTATAGCCGTGAGTGCTGCCAATC  GGTA |

| **BLA-CaM-**  **(mSA2/ binding domain) - His** | ATGGATAAACATTTATTGGCTAAAATTGCTTTATTAGGCGCTGCTCAGCTAGTT ACACTCTCAGCATTTGCTGATCATCCGGAGACATTAGTAAAAGTAAAAGACGCCG AAGATCAACTGGGGGCTCGGGTTGGCTACATTGAGCTTGATCTTAATAGCGGGA AGATACTTGAAAGTTTTCGTCCCGAGGAACGTTTTCCGATGATGTCAACTTTTAA GGTACTGCTTTGTGGTGCCGTTCTGAGCCGTGTAGACGCGGGTCAAGAACAGTT GGGTAGAAGAATTCACTATTCACAGAATGACCTGGTGGAATACTCCCCCGTTACG GAAAAGCATTTGACAGACGGGATGACGGTGCGGGAACTGTGCTCAGCGGCGAT TACTATGAGCGACAACACCGCCGCAAACTTGTTGCTGACGACGATTGGTGGTCC TAAAGAATTAACGGCCTTTCTGCACAATATGGGGGATCACGTCACACGGCTGGA TCGGTGGGAACCGGAGCTGAATGAAGCTATACCTAACGATGAGAGAGACACCA CAATGCCCGTGGCTATGGCGACTACATTACGGAAGCTGTTGACAGGAGAGTTAC TTACACTGGCGTCTAGACAGCAGTTAATCGACTGGATGGAAGCTGATAAGGTAG CCGGGCCATTGTTACGGAGCGCGTTACCTGCGGGGTGGTTTATCGCGGATAAGA GCGGAGCGGGGGAACGGGGTTCTAGAGGGATCATTGCTGCTTTGGGACCGGA CGGCAGCGGAGGGACAGAGGAACAAATCGCCGAATTTAAGGAAGCCTTTTCGT TATTTGACAAGGACGGAGACGGGACTATTACGACGAAGGAGCTTGGCACCGTC ATGCGTTCCCTGGGTCAGAATCCTACAGAAGCCGAGTTACAAGATATGATAAATG AGGTTGACGCTGACGGTAACGGAACTATCGACTTCCCTGAATTTTTAACCATGAT GGCACGCAAAATGAAAGATACAGATAGCGAGGAAGAGATCCGGGAAGCCTTTA GAGTGTTTGACAAAGACGGGAACGGATACATATCAGCCGCTGAGTTGCGGCAC GTTATGACTAACCTTGGCGAAAAACTTACTGACGAAGAAGTAGACGAAATGATA CGTGAGGCTGACATCGACGGAGATGGACAGGTCAACTATGAGGAGTTCGTGCA AATGATGACTGCTGGCGGATCGAGCGGGAAGCCGAGTCGCATCGTGGTAATATA TACAACTGGTTCTCAGGCCACAATGGATGAGCGGAATAGACAGATAGCCGAAAT TGGCGCTAGCTTAATAAAGCACTGGGGCGGAAGTGGCTCCGGTGGTTCAGGTG GCGGTAGCGGTAGCTCCGGAGGAAGTGGCGGATCTGGGGGTGGAGGATCC-(mSA2/BINDING-DOMAIN)—GTCGACAAGCTTGCGGCCGCACTCGAG CACCAC  CACCACCACCAC |
| --- | --- |

**Supplementary Table 3.** Protein sequences used in this study.

| **mSA2** | EFASAEAGITGTWYNQHGSTFTVTAGADGNLTGQYENRAQGTGCQNSPY TLTGRYNGTKLEWRVEWNNSTENCHSRTEWRGQYQGGAEARINTQWN  LTYEGGSGPATEQGQDTFTKVKPSAAS |
| --- | --- |
| **His-FLAG- (mSA2/**  **binding domain)-CaM-BP** | DSHHHHHHGSDYKDDDDKGS - mSA2 / BINDING-DOMAIN -  VDGGSGSGGSGGGSGSSGGSGGSGGGKRRWKKNFIAVSAANR |
| **BLA-CaM-(mSA2/**  **binding domain) - His** | DHPETLVKVKDAEDQLGARVGYIELDLNSGKILESFRPEERFPMMSTFKVL LCGAVLSRVDAGQEQLGRRIHYSQNDLVEYSPVTEKHLTDGMTVRELCSA AITMSDNTAANLLLTTIGGPKELTAFLHNMGDHVTRLDRWEPELNEAIPN DERDTTMPVAMATTLRKLLTGELLTLASRQQLIDWMEADKVAGPLLRSAL PAGWFIADKSGAGERGSRGIIAALGPDGSGGTEEQIAEFKEAFSLFDKDG DGTITTKELGTVMRSLGQNPTEAELQDMINEVDADGNGTIDFPEFLTM MARKMKDTDSEEEIREAFRVFDKDGNGYISAAELRHVMTNLGEKLTDE EVDEMIREADIDGDGQVNYEEFVQMMTAGGSSGKPSRIVVIYTTGSQA  TMDERNRQIAEIGASLIKHWGGSGSGGSGGGSGSSGGSGG SGGGGS - (mSA2 / BINDING-DOMAIN) - VDKLAAALEHHHHHH |
| **His - Avi - 3C - Galectin-3** | DGHHHHHHGSGGLNDIFEAQKIEWHEGSGSGLEVLFQGPGSGGSHMAGPLIVPYNLPLPGGVVPRMLITILGTVKPNANRIALDFQRGNDVAFHFNPRFNENNRRVIVCNTKLDNNWGREERQSVFPFESGKPFKIQVLVEPDHFKVAVNDAHLLQYNHRVKKLNEISKLGISGDIDLTSASYTMI |
| **His - Avi - 3C - S100A7** | DGHHHHHHGSGGLNDIFEAQKIEWHEGSGSGLEVLFQGPGSGSNTQAERSIIGMIDMFHKYTRRDDKIDKPSLLTMMKENFPNFLSACDKKGTNYLADVFEKKDKNEDKKIDFSEFLSLLGDIATDYHKQSHGAAPCSGGSQ |
